# The CRISPR ring nuclease Csx15 oligomerises on cyclic nucleotide binding to regulate antiviral defence

**DOI:** 10.64898/2026.01.21.700848

**Authors:** Haotian Chi, Stephen McMahon, Shirley Graham, Malcolm F White

## Abstract

Prokaryotic type III CRISPR systems signal infection by generating cyclic oligoadenylate (cOA) second messengers, which activate defence proteins allosterically, providing immunity. cOA molecules are typically degraded by extrinsic, stand-alone ring nuclease (RN) enzymes with phosphodiesterase activity, or by the intrinsic RN activity of the effectors themselves. Viruses and plasmids also encode RNs, which can function as anti-CRISPRs (Acr). Eight different families of extrinsic RNs are currently known. Here, we report the structural and biochemical analysis of one of these families: Csx15. We show that Csx15 is a dimeric protein with the ability to bind cyclic tetra-adenylate (cA_4_) molecules in a shared binding site formed by the head-to-tail stacking of dimers in a filament conformation. Some family members are non-enzymatic, relying on the sequestration (sponging) of cA_4_ to regulate the host immune response, while others act as canonical RNs, slowly degrading cA_4_.

## Introduction

CRISPR systems provide adaptive prokaryotic immunity by capturing and storing short segments of DNA from invading mobile genetic elements (MGE) into a CRISPR array in the host genome. This is transcribed and processed into CRISPR RNAs (crRNAs), guiding CRISPR defence systems to target cognate invading nucleic acids during future invasions. Seven major types of CRISPR system have been described [1]. Type III CRISPR systems consist of a multi-subunit interference complex with a Cas10 catalytic subunit. Upon binding to invading RNA, the cyclase domain of Cas10 is allosterically activated, producing cyclic oligoadenylates (cOA) or SAM-AMP as secondary messengers [2-4]. These messengers activate various downstream effectors, initiating diverse immune responses, including nonspecific RNA or DNA cleavage, proteolysis, transcriptional and translational regulation, and even membrane disruption (reviewed in [5]). The activation of these effectors can cause cell growth arrest or death, thus providing protection at the population level [6].

In reality, many type III CRISPR systems include a specialised class of enzymes, collectively known as “ring nucleases” (RN) that degrade the cyclic nucleotide, providing a mechanism to deactivate the immune response and avoid unnecessary cell death. The first CRISPR-associated ring nuclease (Crn1) was identified biochemically [7], followed quickly by Crn2 and Crn3 [8, 9]. These enzymes are all specific for degradation of cA_4_, the most commonly used signalling molecule in type III CRISPR systems [10]. Recently, a further enzyme, Crn4, was described which has a much broader substrate specificity, degrading all cOA species tested [11]. In addition, many of the effector proteins activated by cOA species have intrinsic RN activity and are therefore self-limiting [12-19]. A recent bioinformatic analyses revealed that the ring nucleases are diverse and abundant in type III CRISPR systems, adding two more cA_4_ specific enzymes (Csx16 and Csx20) to the growing list [20]. Viruses also encode ring nucleases in their genomes, where they function as anti-CRISPR (Acr) proteins by intercepting and destroying cOA second messengers [8, 21].

Csx15 was originally proposed as a candidate RN based on its amino acid sequence and genomic neighbourhood analysis [22]. Analysis of ∼1000 type III CRISPR loci in 40,000 complete prokaryotic genomes revealed that Csx15 was present in 18 loci, often co-occurring with Crn1 [20]. About 25% of *csx15* genes in the dataset were orphans, not associated with a CRISPR locus. Alphafold3 (AF3) [23] modelling predicted a CARF-like Rossmann fold for Csx15. A representative example from *Pseudomonas fluorescens* (PfCsx15) was tested for RN activity but found to have little or no activity *in vitro* [20]. However, PfCsx15 is an orphan protein and could thus have diverged in function.

Here, we investigate two Csx15 orthologues, the orphan PfCsx15 and the CRISPR-associated *Chlorobaculum limnaeum* Csx15 (CliCsx15). PfCsx15 specifically sequesters cA_4_ to inhibit the cA_4_-activated ribonuclease activity of a CRISPR effector protein. CliCsx15 exhibits cA_4_-specific RN activity *in vitro* and can neutralise CRISPR defence *in vivo*. Both proteins adopt higher oligomerisation states induced by cA_4_ binding. Crystal structures of PfCsx15 and CliCsx15 reveal a conserved dimeric structure and the molecular basis for multimerization in a head-to-tail conformation that sandwiches cA_4_ between dimers. The likely functions of these unusual cA_4_ binding proteins is discussed.

## Results

### PfCsx15 specifically inhibit cA_4_-activated CRISPR effectors

Although PfCsx15 was previously identified as a candidate RN, very limited nuclease activity was observed using HPLC analysis [20]. To further investigate the function of PfCsx15, we tested whether the protein could inhibit a range of type III CRISPR effectors *in vitro*. We examined a cA_3_-activated DNase *Vibrio metoecus* (Vme) NucC [24], a cA_4_-activated ribonuclease TTHB144 from *Thermus thermophilus* HB8 [12] and the cA_6_-activated ribonuclease *Mycobacterium tuberculosis* (Mtb) Csm6 [25] using established fluorescent assays. We observed a progressive inhibition of TTHB144 in the presence of high concentrations of PfCsx15. At three concentrations of cA_4_ (1, 2 or 4 M), preincubation with PfCsx15 for 15 min led to a reduction in TTHB144 ribonuclease activity when the molar ratio of PfCsx15 dimer to cA_4_ concentration was higher than approximately 0.6 (Figure 1). Weaker inhibition was observed for VmeNucC (Supplementary Figure 1) and MtbCsm6 (Supplementary Figure 2). These data suggest that PfCsx15 is specific for cA_4_, and does not primarily function as a RN, but rather exhibits properties more characteristic of a cyclic nucleotide binding, or sponge, protein, analogous to the anti-CBASS protein Acb2 [26].

**Figure 1.**
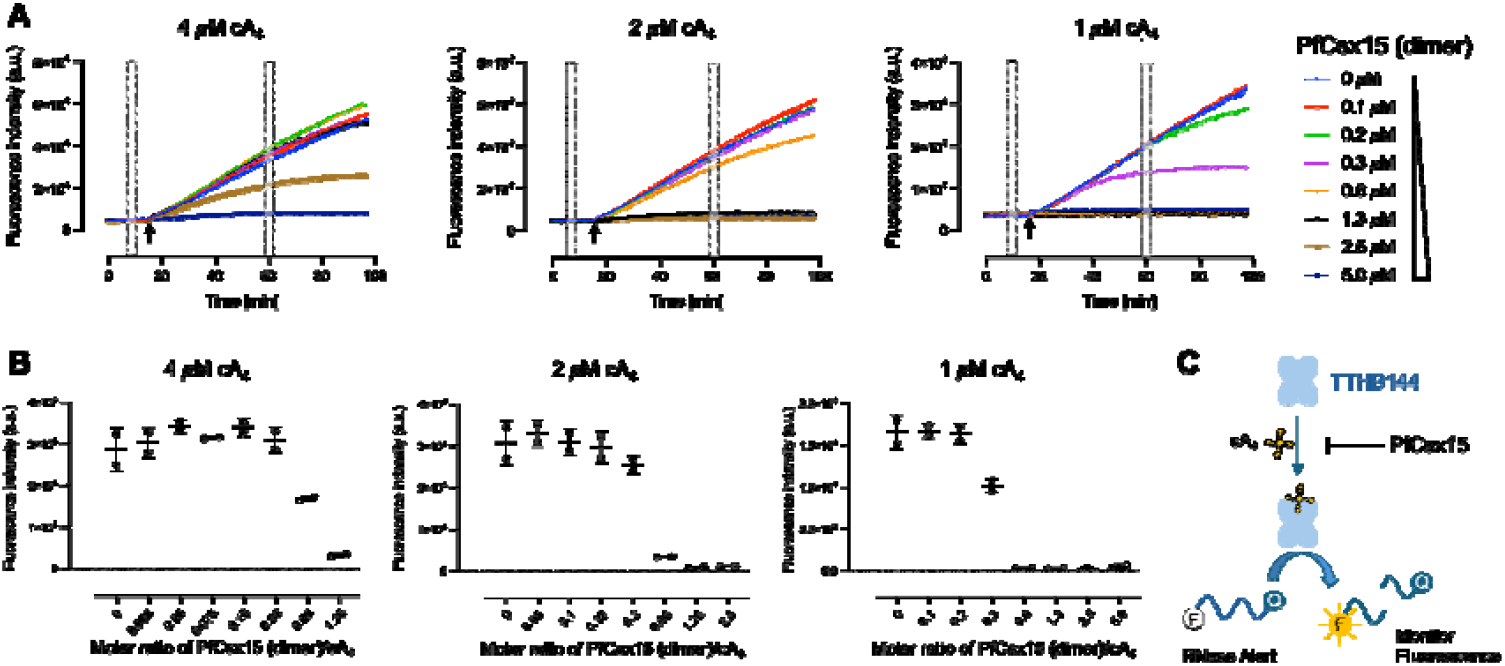
PfCsx15 specifically inactivates cA_4_-induced RNase TTHB144. **A**. Plot of baseline-corrected fluorescence signal in **B** against the ratio of Csx15 (dimer) to cA_4_. The average fluorescence signal in **B** from 59-61 min was corrected using the average baseline signal from 9-10 min. Data of two replicates were presented as mean ± s.d. The RNase activity of TTHB144 was inhibited when the ratio of Csx15 (dimer) to cA_4_ exceeded 0.6. **B**. RNase alert assay with detection of fluorescence signal emitted by RNA cleavage when RNases were presented. RNaseAlert substrates (100 nM) were incubated with the Csx15 (0-5 uM dimer) and cA_4_ (1, 2 and 4 µM, respectively) at 35 °C for 15 min, before adding RNase TTHB144 (500 nM dimer). The fluorescence signal was plotted against time. **C**. Schematic representation of the fluorogenic biochemical assay.

### PfCsx15 oligomerises on cA_4_ binding

Oligomeric states of PfCsx15 were examined using dynamic light scattering (DLS) in the presence or absence of different cOA species. Addition of cA_4_, but not cA_3_ or cA_6_, to PfCsx15 resulted in a significant increase in global particle size (Figure 2A). Additionally, size exclusion chromatography (SEC) showed that PfCsx15 eluted with a retention time consistent with a dimer in the absence of cA_4_, while a significant shift to a shorter retention time (and thus larger size) was observed in the presence of cA_4_ (Figure 2B). These observations suggested that cA_4_ induces PfCsx15 to form high molecular weight complexes.

**Figure 2.**
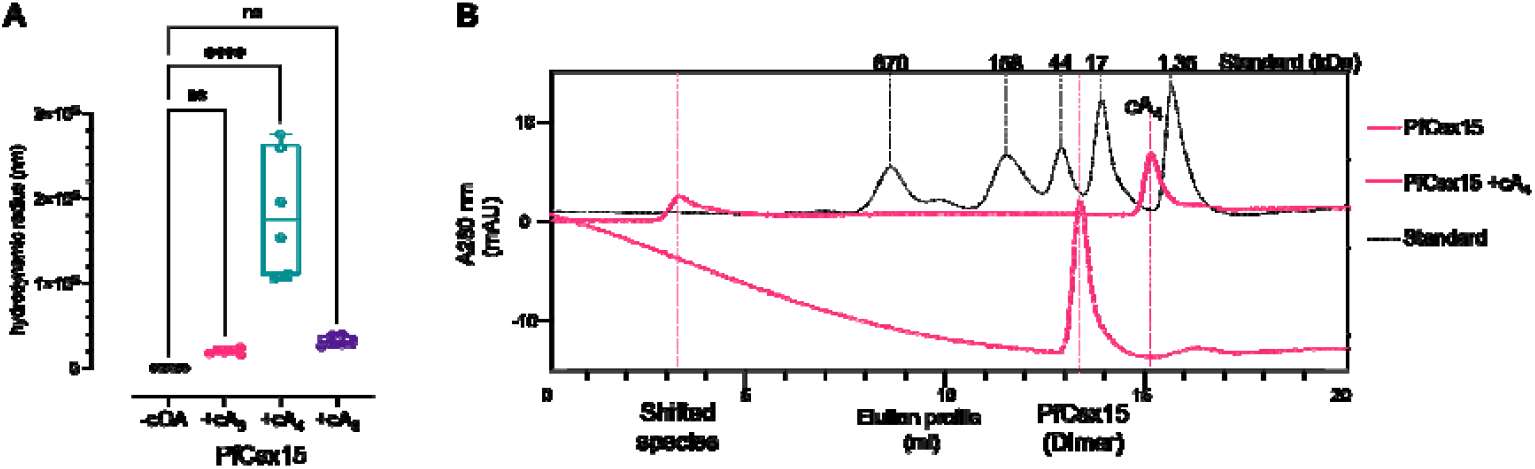
cA_4_ induces the oligomerization of PfCsx15. **A**. Dynamic Light Scattering (DLS) of Csx15 in the presence or absence of a range of cOAs. Csx15 assembles into significant larger molecular species only in the presence of cA_4_. Statistical analysis was preformed using two-way ANOVA multiple comparisons, followed by Šídák’s multiple comparisons test (\*\*\*\**P*<0.0001). **B**. Size exclusion chromatography (SEC) of Csx15 with and without cA_4_. A molar ratio of Csx15 (dimer) to cA_4_ is 2:1. Gel filtration standard (Bio-Rad) includes thyroglobulin (670,000), g-globulin (158,000), ovalbumin (44,000), myoglobin (17,000), and vitamin B12 (1,350).

### Crystal structure of PfCsx15

To better understand the structural organisation of PfCsx15, we crystallised the protein and solved the structure at a resolution of 1.88 Å (Supplementary Table 1). The dimeric structure corresponds closely to that predicted by AF3 previously [20], comprising a Rossmannoid fold with a central 5-stranded β-sheet flanked by α-helices. The crystal packing positions dimers in a repeating head-to-tail conformation, generating a potential shared binding site between dimers (Figure 3A). Two β-hairpins arising from the top surface of each dimer form interactions with the dimer positioned above in the crystal lattice. In the dimer:dimer interface, we observed two phosphate ions, which might indicate the binding site of cA_4_. The top dimer (orange) contributes a number of conserved arginine residues (R116, R120, R123), which interact with the phosphate ions, along with conserved H82 (Figure 3B). The bottom dimer (teal) contributes the conserved residues S9 and H11. The 6 arginine residues appear suitably positioned to interact with the phosphate groups in bound nucleotide ligands. Unfortunately, we were unable to co-crystallise the protein with cA_4_, as the protein came out of solution when the nucleotide was added. A selection of Csx15 orthologues is shown aligned in Supplementary Figure 3.

**Figure 3.**
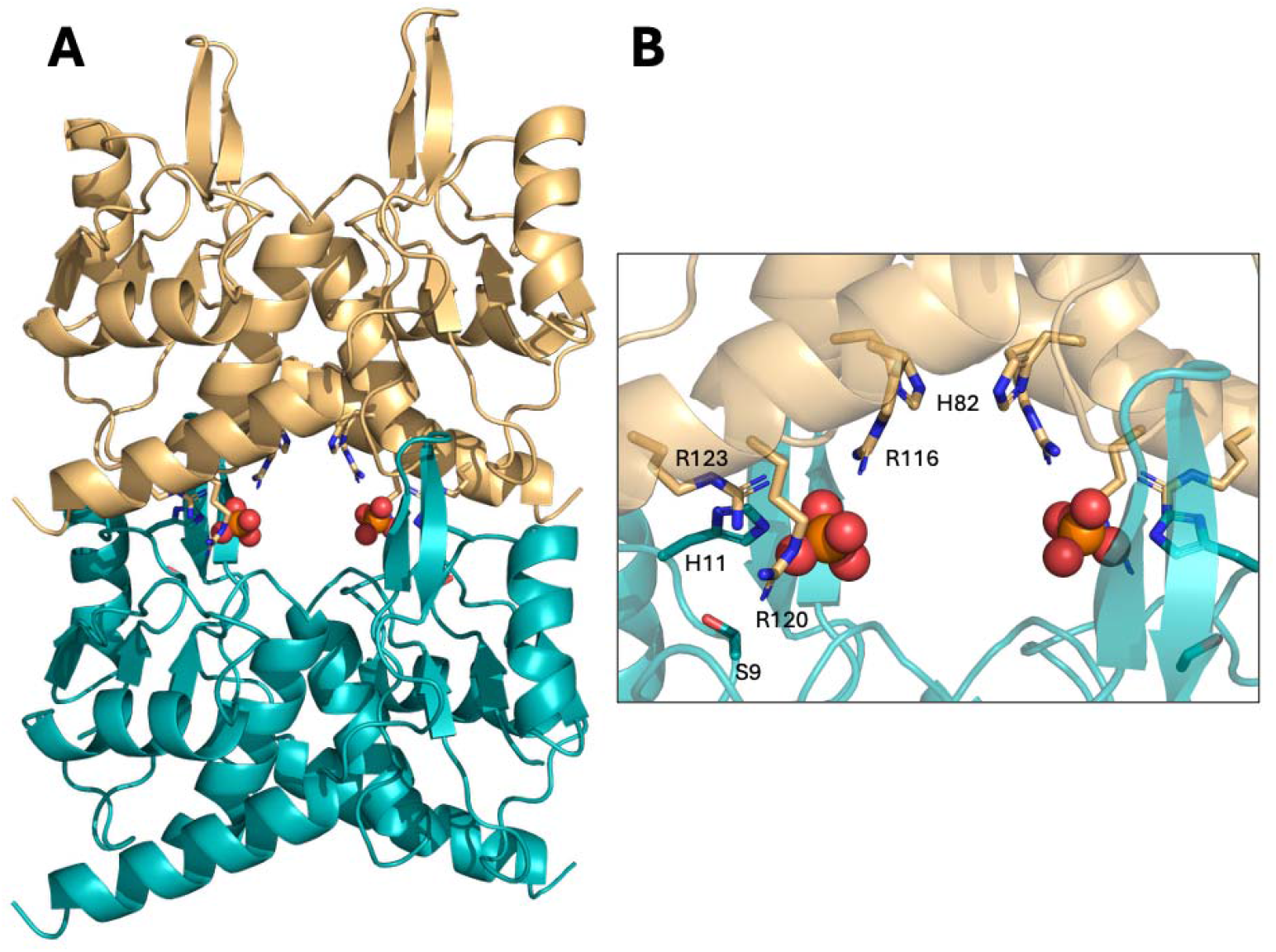
Crystal structure of PfCsx15. **A**. Crystal structure of PfCsx15, showing dimer:dimer stacking in the crystal lattice with a head-to-tail organisation. **B**. Close-up view of the dimer:dimer interface, showing conserved residues H82, R116, R120 and R123 in the top dimer, S9 and H11 in the bottom dimer. The positions of two phosphate ions present in the crystal structure are shown as spheres.

### Mutation of conserved interface residues reduces filament formation

To test for an interaction with cA_4_, residues H11 (bottom dimer) and R123 (top dimer) of PfCsx15 were independently mutated to alanine. DLS assays were first performed to examine the oligomeric states of PfCsx15 wild type and variants (Figure 4A). The formation of high molecular weight species upon addition of cA_4_ was largely abolished for the H11A variant (Figure 4A). Although a higher oligomeric state was still observed for the R123A variant, the global particle size was significantly less than that observed for the wild type. These findings suggest that both residues play a role in the formation of higher-order oligomeric states and reinforce the conclusion that the dimer:dimer interface is the site of cA_4_ binding.

**Figure 4.**
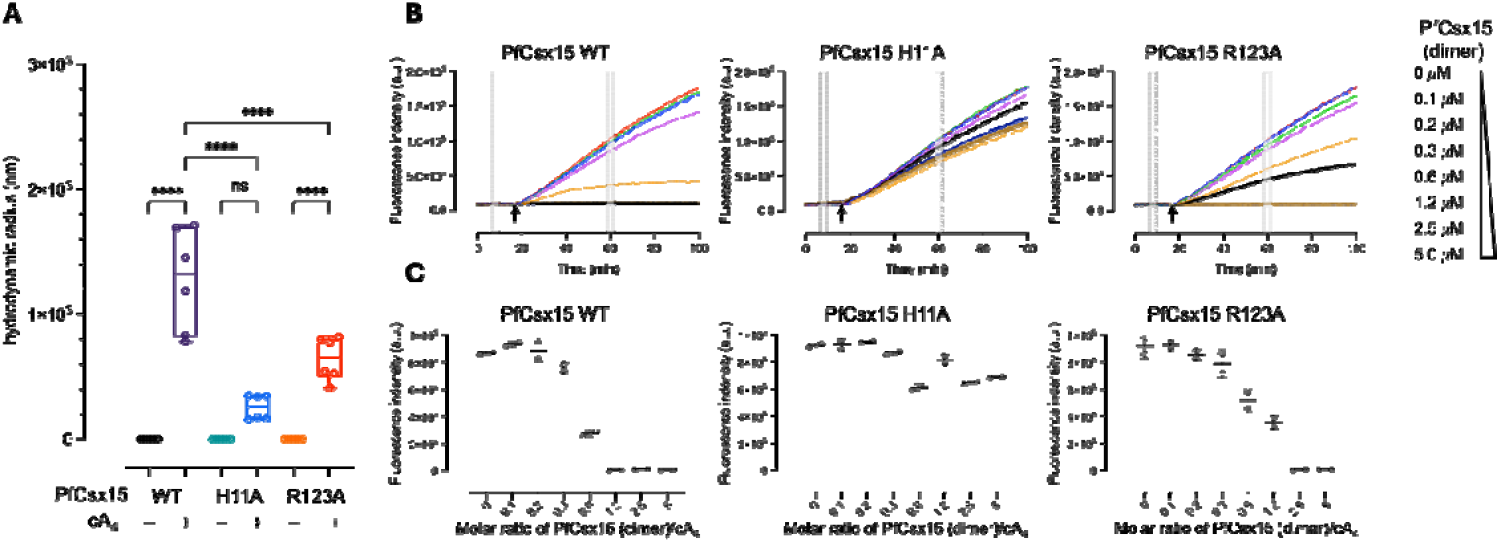
Characterisation of PfCsx15 variants H11A and R123A. **A**. Dynamic Light Scattering (DLS) of PfCsx15 wild type and variants. Mutation of conserved residues H11 and R123 abolished the cA_4_-induced multimerization of PfuCsx15. Statistical analysis was preformed using two-way ANOVA multiple comparisons, followed by Turkey’s multiple comparisons test (\*\*\*\**P*<0.0001, ^ns^*P* =0.0726). **B**. Fluorescent assay of PfCsx15 wild type and variants. RNaseAlert substrates (100 nM) were incubated with the Csx15 (0-5 uM dimer) and cA_4_ (1.1 µM) at 35 °C for 15 min, before adding RNase TTHB144 (500 nM dimer). The fluorescence signal was plotted against time. **C**. Plot of baseline-corrected fluorescence signal in **B**. Data of two replicates were presented as mean ± s.d. There was no observed inhibition of cA_4_-induced TTHB144 ribonuclease activity when H11A was present.

Ribonuclease inhibition assays were subsequently carried out to investigate the ability of the variant proteins to sequester cA_4_ (Figure 4C). No inhibition of cA_4_-activated TTHB144 ribonuclease activity was observed, even at very high concentrations of the H11A variant (Figure 4B). For the R123A variant, full inhibition occurred at higher protein-to-cA_4_ ratios than that required for the wild-type protein (Figure 4B). Collectively, these observations indicate that H11 plays a major role in cA_4_ binding and multimerisation while R123 plays a secondary role.

### CliCsx15, a type III CRISPR associated Csx15, is a cA_4_-specific ring nuclease

As PfCsx15 is encoded by an orphan gene which is not linked to a type III CRISPR locus, it may have diverged in function. We therefore selected and cloned a gene encoding Csx15 from the type III CRISPR locus of *Chlorobaculum limnaeum* strain DSM 1677, hereafter referred to as CliCsx15 (Figure 5A). The protein was expressed and purified to homogeneity (Supplementary Figure 4). The RN activity was tested by incubating cOAs (fixed at 80 µM) with increasing concentrations of CliCsx15 for 60 min at 30 °C, followed by HPLC analysis. Linear products A_2_-p and A-p were observed at the higher protein concentrations (Figure 5B). The observation of 3’-phosphate termini (rather than 2’,3’-cyclic phosphate termini) on the products suggests that this is a hydrolytic reaction, involving a water nucleophile. This contrasts with ring nucleases such as AcrIII-1, which degrade cA_4_ using in-line attack of the adjacent 2’-hydroxl group on the substrate [8]. The CliCsx15 RN activity was unaffected by the presence or absence of EDTA/Mg^2+^/Mn^2+^, suggesting that activation of the water nucleophile is not mediated by a divalent metal ion (Supplementary Figure 5). The turnover rate was very slow, with only ∼50% of cA_4_ degraded over 60 min at an enzyme:substrate ratio of 1:8. No RN activity was detected for cA_3_ or cA_6_ substrates (Supplementary Figure 6).

**Figure 5.**
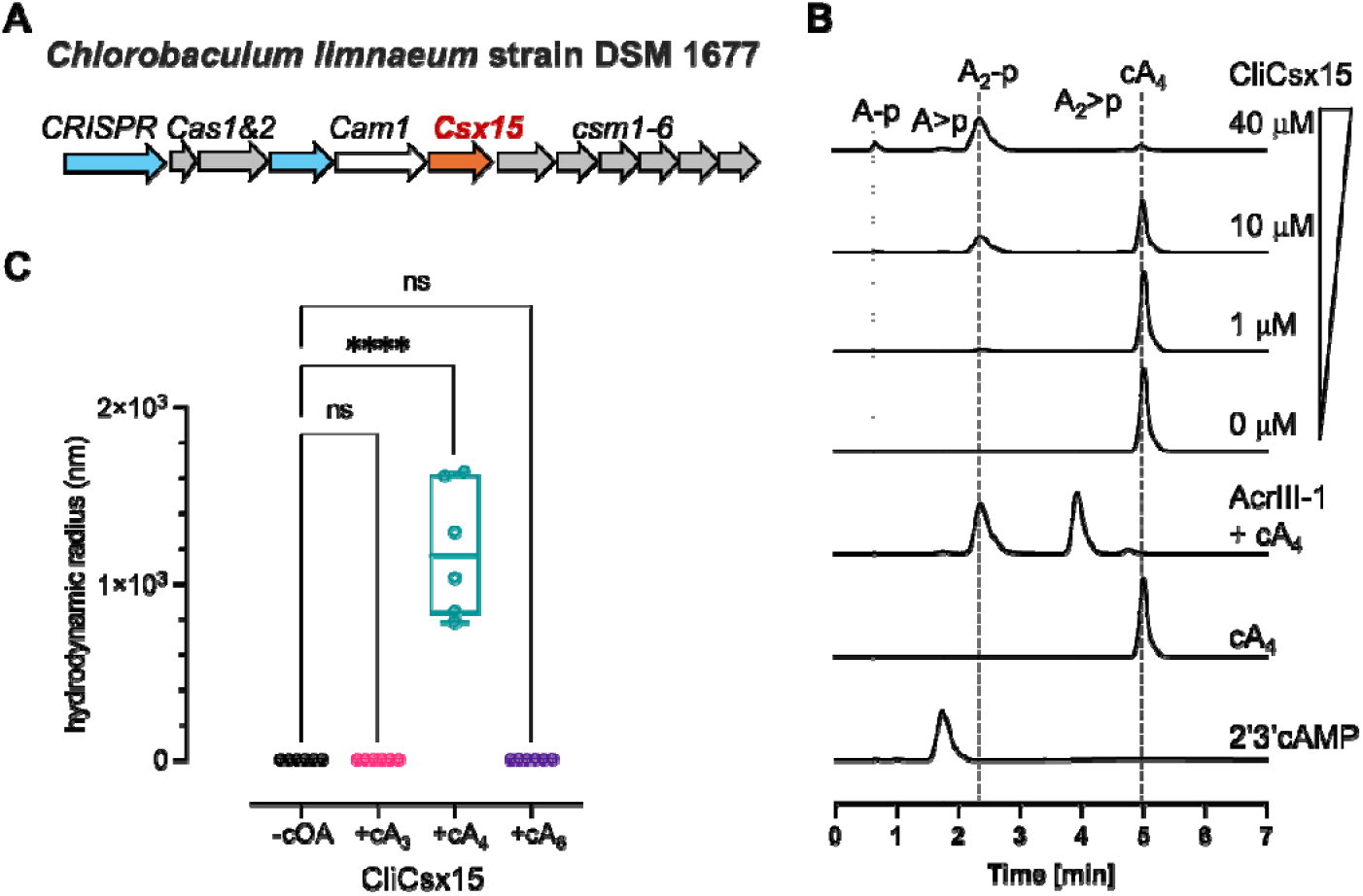
CliCsx15 is a cA_4_-specific ring nuclease. **A**. Genomic context of the Csx15-associated type III-A CRISPR system in *C. limnaeum* strain DSM 1677. **B**. Ring nuclease activity of CliCsx15. cA_4_ (80 M) incubated with CliCsx15 (0, 1, 10 and 40 M) for 60 min at 30 °C was analysed using HPLC. **C**. DLS analysis. cA_4_ promotes the increase in global particle size. Statistical analysis was preformed using two-way ANOVA multiple comparisons, followed by Šídák’s multiple comparisons test (\*\*\*\**P*<0.0001).

We also tested whether cA_4_ promoted the oligomerisation of CliCsx15. DLS was used to examine oligomeric states in the presence or absence of different cOAs. A significant increase in particle size was observed only upon cA_4_ addition to CliCsx15, consistent with the observations for PfCsx15 (Figure 5C).

### Crystal Structure of CliCsx15

We crystallised the apo form of CliCsx15, solving the structure at the extremely high resolution of 0.9 Å (Figure 6A). A sample of the electron density for residues at the interface is shown in Supplementary Figure 7. Overall the dimeric structures of PfCsx15 and CliClx15 are similar, with an RMSD of 3.0 Å over 208 residues. Notably, the same head-to-tail packing was observed in both structures, strengthening the hypothesis that this might represent the biologically relevant interface. To investigate this further, we generated a model of four monomers of CliCsx15 along with 4 AMP molecules (analogous to cA_4_) using AF3. The AF3 model (Figure 6B), predicted with high confidence (ipTM 0.81; pTM = 0.84), matched very well to the organisation of CliCsx15 dimers observed in the crystal lattice. The individual dimers of the crystal structure and AF3 model align with a RMSD of 0.81 Å over 256 residues. The four AMP molecules adopt orientations sandwiched in the dimer:dimer interface (Figure 6C, D), in suitable positions to interact with the conserved arginine residues contributed by the top dimer. Conserved residues H13 and Y113, equivalent to H11 and F106 in PfCsx15 (Supplementary Figure 8), flank the adenosine base of AMP at two positions in the dimer:dimer interface (Figure 6D). Conserved residue H90 sits in a central position in the top dimer, suitably located to participate in catalysis.

**Figure 6.**
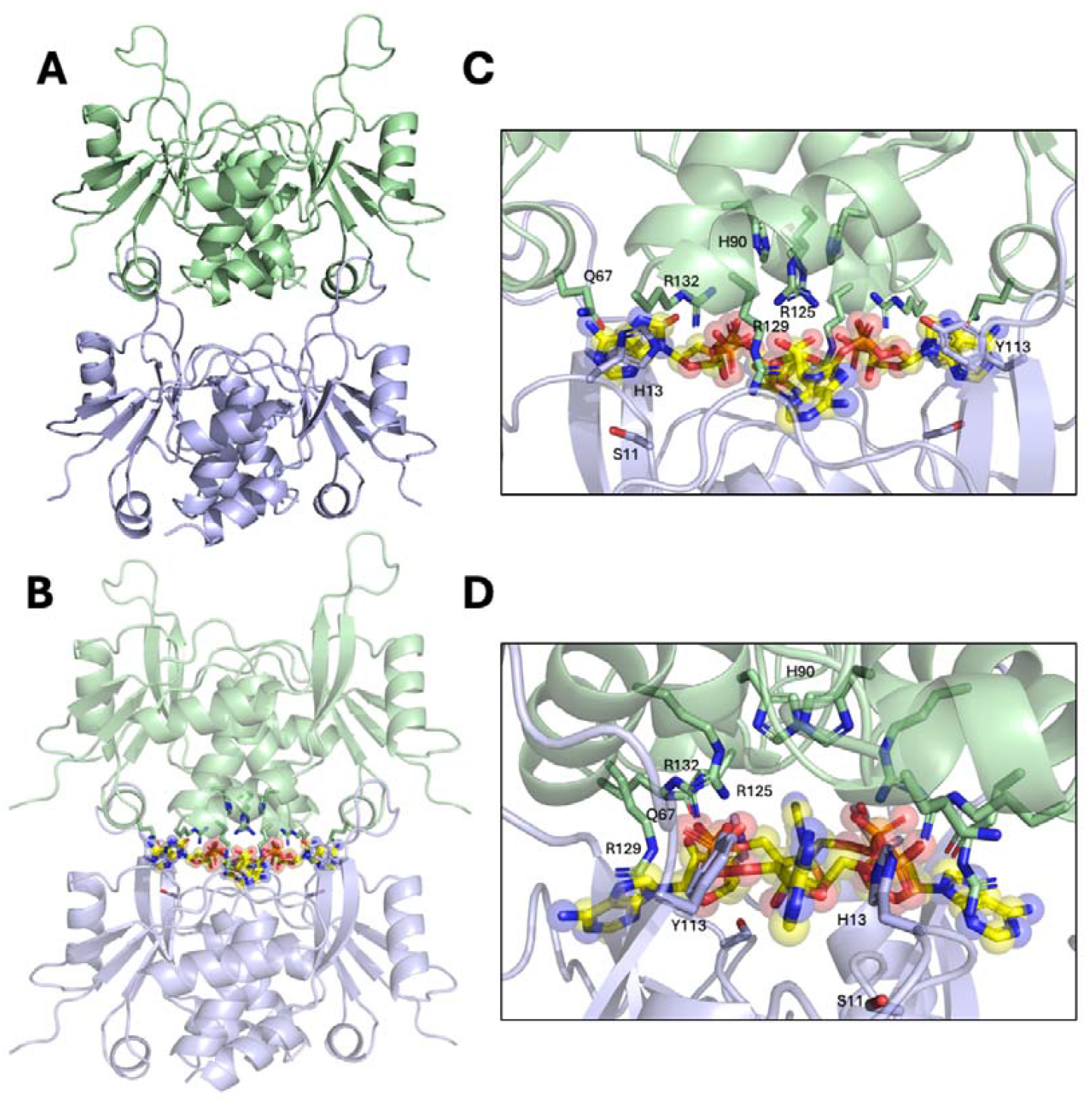
Crystal structure and AF3 modelling of CliCsx15. A. Crystal structure of CliCsx15, showing dimer:dimer stacking in the crystal lattice with a head-to-tail organisation, similar to PfCsx15. B. AF3 model of CliCsx15 together with 4 AMP ligands. The AMP molecules cluster in the dimer:dimer interface, suggesting the expected binding site for cA_4_. C. Close-up view of the dimer:dimer interface in the AF3 model, showing the positions of conserved residues (labelled) D. Orthogonal view of the dimer:dimer interface.

To test the functional roles of H13 and H90, we mutated each residue to alanine and examined the ability of the variants to multimerise on cA_4_ binding and to degrade the cyclic nucleotide. The RN activity of H13A was abolished, while only very low RN activity was observed for the H90A variant (Figure 7A), implicating both residues in the catalytic mechanism. The cA_4_-induced oligomerisation was abolished when the residue H90 was mutated, whereas variant H13A had subtle effects on its oligomeric state (Figure 7B).

**Figure 7.**
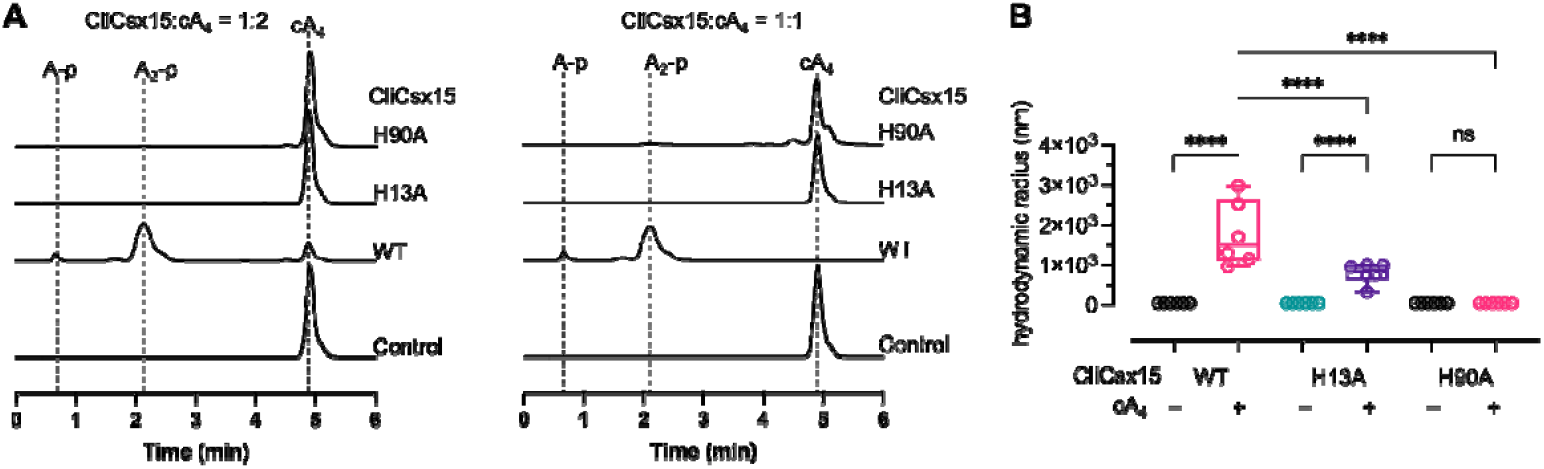
Ring nuclease activity and oligomeric states of CliCsx15 variants. **A**. RN activity of CliCsx15 wild-type and variants. cA_4_ (50 M) incubated with CliCsx15 wt and variants (25 or 50 M dimer) for 60 min at 30 °C was analysed using HPLC. No ring nuclease activity was detected in the presence of variant H13A, whereas limited cleavage activity was observed in the high concentration of variant H90A. **B**. DLS analysis. cA_4_-induced multimerization was completely abolished when a conserved residue H90 was mutated, while the mutation of H13 had less effects on its oligomerisation state. Statistical analysis was preformed using two-way ANOVA multiple comparisons, followed by Šídák’s multiple comparisons test (\*\*\*\**P*<0.0001).

Taken together, our biochemical, X-ray structural and modelling studies provide a strong prediction that Csx15 adopts a linear head-to-tail multimeric structure when bound to cA_4_, reminiscent of the ring nuclease Csx3 [9].

### CliCsx15 can neutralise CRISPR defence *in vivo*

To assess the *in vivo* function of Csx15, we tested CliCsx15 in a well-established recombinant type III-A CRISPR system from *Mycobacterium tuberculosis* (MtbCsm), expressed in *E. coli*. This system generates a range of cOA species and activates the cA_4_-dependent effector *Thioalkalivibrio sulfidiphilus* Csx1 to confer plasmid immunity. *E. coli* C43 cells expressing MtbCsm were transformed with a pRATDuet plasmid encoding Csx1 alone or in combination with CliCsx15, together with a tetracycline resistance gene (*tetR*). When Csx1 was activated by cA_4_ produced by the MtbCsm complex programmed with a CRISPR RNA (crRNA) targeting *tetR* transcripts, a reduction in transformation efficiency was observed, as anticipated. Co-expression with CliCsx15 restored cfu to near-control levels (Figure 8A, B). As a negative control, MtbCsm did not produce cA_4_ when programmed with a crRNA targeting the pUC plasmid and no reduction in transformants was observed (Figure 8).

The neutralisation of cA_4_-mediated plasmid immunity suggested that CliCsx15 effectively reduced the intracellular cA_4_ concentration.

**Figure 8.**
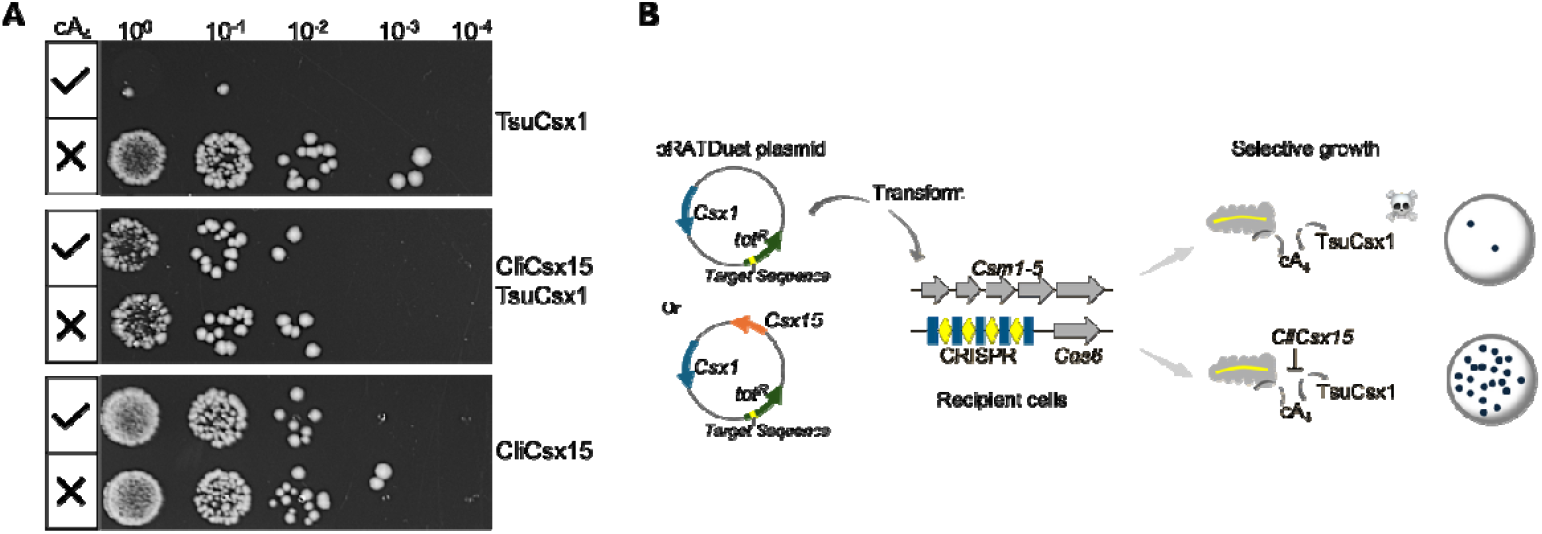
CliCsx15 relieves cA_4_-mediated plasmid immunity. **A**. Plasmid challenge assay. Serial dilutions of *E. coli* transformants expressing the MtbCsm system together with effector Csx1, ring nuclease CliCsx15, or both were spotted onto antibiotic-selective agar plates with inducer to activate expression of all plasmids. Coexpression of CliCsx15 restored transformation efficiency in cells with cA_4_-activated TsuCsx1. Two biological replicates, each with two technical replicates, were performed. Representative plates are shown. **B**. Schematic representation of the experimental system.

## Discussion

Cyclic nucleotide second messengers are a key feature of both eukaryotic and prokaryotic antiviral immunity, providing a convenient means to amplify the primary signal of infection. Typically, these cyclic nucleotides bind and allosterically activate defensive effector proteins to elicit an immune response. An unusual feature of type III CRISPR systems is the near-ubiquity of a mechanism to degrade these cyclic nucleotides via intrinsic or extrinsic ring nucleases, providing a means to swich off this defence pathway when it is no longer needed. Six families of extrinsic cA_4_-specific RNs have been described [20, 21], in addition to an unusual RN (Crn4) which is capable of degrading all tested cOA species [11]. Here, we tested the enigmatic candidate RN Csx15, demonstrating that it binds specifically to cA_4_, forming a head-to-tail, extended filament conformation. Some family members are capable of slowly degrading cA_4_ while others appear to sequester (sponge) the cyclic nucleotide.

Sponging of cyclic nucleotides is commonly encountered as an anti-defence strategy to combat the CBASS and Thoeris immune systems in bacteria [26-29]. This appears effective when the levels of second messenger produced are quite low. In response, some bacteria encode the Panoptes system, which constitutively generates “decoy” cyclic nucleotides to keep a membrane-bound effector deactivated [30, 31]. When a phage with a sponge invades cells with Panoptes, sequestration of the decoy molecules results in the activation of Panoptes defence. In contrast to these pathways, type III CRISPR systems typically make much larger quantities of second messenger molecules [24, 32, 33], which could make sponging a fruitless approach. In response, viruses encode RNs and phosphodiesterases to degrade these molecules [34].

What then is the rationale for Csx15 – a protein that is a good cA_4_ sponge but a very slow ring nuclease? Firstly, we note that Csx15 was not observed in any phage genomes [20], suggesting that it may not have utility as an anti-CRISPR. We have observed effective neutralisation of a cA_4_-activated CRISPR effector by Csx15 *in vivo* (Figure 6), but it should be borne in mind that this involves over-expression of the protein from a multicopy plasmid, which could enhance its effectiveness. Perhaps, when part of a CRISPR system, Csx15 serves to mop up any small amounts of cA_4_ that are generated aberrantly, for example by binding of a cellular transcript with a partial match to the crRNA of the type III system. Csx15 could thus act as a dampener to reduce auto-immunity while avoiding the unwanted neutralisation of the system when fully activated by phage infection.

## Methods

### Cloning

Synthetic genes encoding PfCsx15 (WP_191946534.1) and CliCsx15 (WP_069809204.1), purchased from Integrated DNA Technologies (IDT)) were codon optimised for expression in *Escherichia coli* and constructed into the pEHisV5TEV vector between the *Nco*I and *BamH*I restriction sites. Mutants were created by the site-directed mutagenesis using primers with the desired mutations. The *E. coli* DH5 α strain was used for the cloning and mutagenesis. Sequence integrity and success mutations were confirmed by sequencing (Eurofins Genomics). All synthetic genes and primers used in this study were listed in supplementary table 2.

For the plasmid challenge assay, the *CliCsx15* gene was cloned into the multiple cloning site - 1 (MCS-1) between the *Nde*I and *Xho*I restriction sites of pRATDuet, under the control of a T7 promoter. For co-expression, the *Thioalkalivibrio sulfidiphilus* (*Tsu) Csx1* gene was cloned into the MCS-2 of pRATDuet-CliCsx15 plasmid between *Nco*I and *Sal*I sites, under the control of a pBAD promoter.

### Protein expression and Purification

The expression and purification of PfCsx15 was previously described [20], the same protocols were used for CliCsx15. In brief, *E. coli* C43 (DE3) strain was used for the protein expression. 2 l of cell culture were induced with 0.4 mM isopropyl-β-D-1-thiogalactoside (IPTG) when OD_600_ reached 0.6-0.8 and grown overnight at 25°C. Cell pellets were harvested by centrifugation and lysed by sonication for purification. Proteins were purified using immobilised metal affinity chromatography (IMAC) and size exclusion chromatography. The his-tag was removed by tobacco etch virus (TEV) protease. The identity and purity of protein were confirmed using SDS-PAGE and pure proteins were stored frozen at -70°C. For the determination of molarity, Csx15 was considered to be dimeric.

### Ring nuclease assay

Ring nuclease activity was examined by incubating enzymes (1, 10 and 40 *μ*M) with 70 *μ*M cOA species (Biolog) in the reaction buffer of 20 mM Tris-HCl, 250 mM NaCl, pH7.5 at 30°C for 60 min. For multiple turnovers, 2 *μ*M CliCsx15 (dimer) was incubated with 10-fold excess cA_4_ at 30°C for the time points of 5, 10, 30, 60, 120 and 240 min. For single turnover, 40 *μ*M CliCsx15 (dimer) was incubated with 25 *μ*M cA_4_ at 30°C for the time points of 1, 3, 5, 10, 15, 30, 45 and 60 min. The reactions were terminated by adding two equivalent volumes of cold methanol and vortex. Samples were vacuum dried, before resuspension in water for HPLC analysis.

### HPLC analysis

HPLC was conducted on a Ultimate3000 UHPLC system (Thermo Fisher Scientific) with a C18 column (Kinetex EVO 2.1 × 50 mm, particle size 2.6 *μ*m). The column temperature was set at 40°C and the absorbance was monitored at 260 nm. Samples were analysed by gradient elution with solvent A (20 mM ammonium acetate, pH8.5) and solvent B (methanol) at a flow rate of 0.3 ml/min as follows 0-0.5 min, 1% B; 0.5-6 min, 1-15% B; 6-7 min, 100% B.

### Dynamic light scattering (DLS)

DLS measurements were conducted on a Zetasizer Nano S90 (Malvern) instrument to evaluate the hydrodynamic radii of Csx15 and CliCsx15 in the presence or the absence of cOAs. 1 mg/ml proteins were mixed with or without 70 μM cOAs in the buffer of 20 mM Tris-HCl, pH 7.5, 150 mM NaCl and 10% glycerol. Samples were centrifuged at 14,000 rpm for 10 min prior to measurement. The experiment was carried out at 4°C with 3 measurements of 13 runs in two independent experiments. Statistical analysis was preformed using two-way ANOVA multiple comparisons, followed by Šídák’s multiple comparisons test (Prism, version 10.2.2)

### Fluorogenic biochemical assay

Assays were carried out on a fluorescence plate reader (FLUOstar Omega, BMG Labtech) with a Greiner 96 half-area plate. Csx15 (0, 0.1, 0.2, 0.3, 0.6, 1.2, 2.5 and 5 μM dimer) was incubated with cA_4_ (1, 2 and 4 μM) in the presence of RNAse-Alert (IDT) substrate (100 nM). Reactions were incubated at 35 °C for 15 min in a buffer of 20 mM Tris-HCl, 100 mM NaCl, pH 7.5, before adding *Thermus thermophilus* HB8 cA_4_ activated ribonuclease TTHB144 (500 nM dimer). Fluorescence intensity was monitored up to 90 min with 30 s intervals (λ_ex/em_ 485 / 520 nm). Csx15 with the same gradient was incubated with cA_3_ (2 and 5 μM) in the presence of FAM: Iowa Black® double-stranded DNA substrate (100 nM) in a buffer of 20 mM Tris-HCl, pH7.5, 250 mM NaCl and 10 mM MgCl2. Reactions were conducted at 35°C for 15 min, before adding cA_3_ activated DNase *Vibrio metoecus* NucC (250 nM trimer) and continued for 85 min with same fluorescence measurements. For cA_6_ activated ribonuclease *Mycobacterium tuberculosis* Csm6, Csx15 was incubated with cA_6_ in the same ratio of Csx15/cA_3_ and RNase-Alert in a buffer of 20 mM Tris-HCl, pH7.5, 250 mM NaCl and 100 mM potassium L-glutamate with the same fluorescence measurements. The curves are the mean of replicates (Prism, version 10.2.2).

### Protein crystallisation

Immediately prior to crystallisation both proteins, PfCsx15 at 15 mg mL^-1^ and CliCsx15 at 16 mg mL^-1^, were centrifuged at 14000g. Sitting drop vapour diffusion experiments were set up at the nanoliter scale at 1:1 and 2:1 protein to reservoir solution ratios. Four commercially available crystallisation screens were screened, upon completion plates were sealed and incubated at 293 K. Of the many conditions that yielded crystals, PfCsx15 crystals used in data collection were grown from 22.5% PEG smear medium, 10% glycerol, 0.2M sodium potassium phosphate pH 7.5, and 0.1M HEPES pH7.5. CliCsx15 crystals used in data collection were grown from 22% PEG smear broad, and 0.1M Tris pH 8.8. All crystals were cryoprotected with 25% glycerol prior to harvesting and cryo-cooling in liquid nitrogen.

### X-ray data collection, structure solution, and refinement

All X-ray data were collected at the Diamond Light Source. Data for PfCsx15 were collected at wavelength 0.9537 Å, 100 K, on beamline I04 to 1.88 Å, whereas data from CliCsx15 crystals were collected on IO3 at 0.7918 Å to 0.90 Å resolution. All data were automatically processed using Xia2 [35]. The structures were solved by phasing the data using PhaserMR [36] in the CCP4 suite [37], using models generated by AlphaFold 3 [23]. Initial B-factors were modelled in Phenix [38]. Model refinement was achieved by iterative cycles of REFMAC5 [39], all hydrogens generated but not written to the output file, with manual model manipulation in COOT [40]. Anisotropic B factor refinement was performed in the case of CliCsx15 and isotropic for PfCsx15.

As Pfcsx15 was crystallised in the presence of Sodium / Potassium phosphate, a phosphate ion was modelled in the structure. The geometric qualities of each structure were monitored throughout using Molprobity [41]. Data and refinement statistics are shown in Supplementary Table 1. Final coordinates have been validated and deposited in the Protein Data Bank with deposition codes 9TEU for CliCsx15, and 9TET for PfCsx15 along with the original data.

### Analytical size exclusion chromatography

To analyse the oligomeric state of Csx15, 100 μl protein (1 mg/ml) with or without cA_4_ was analysed using a Superose 6 Increase 10/300 GL size-exclusion column, which was equilibrated in 20 mM Tris-HCl, pH 8.0, 250 mM NaCl, 10% glycerol and 1 mM DTT. The eluted fractions were then analysed by SDS-PAGE gel (Invitrogen). Standards (catalog number 1511901, BioRad) were analysed in the same running condition, comprising thyroglobulin (670,000 Da), γ-globulin (158,000 Da), ovalbumin (44,000 Da), myoglobin (17,000 Da), and vitamin B12 (1,350 Da).

### Plasmid challenge assays

Plasmids used in this study for expression of the programmed type III MtbCsm system include pCsm1-5 (containing Csm interference genes *cas10, csm3, csm4* and *csm5* from *M. tuberculosis* and *csm2* from *M. canettii*), pCRISPR-TetR and pCRISPR-pUC (containing *M. tuberculosis cas6* and a CRISPR array targeting a tetracycline-resistance gene or a pUC multiple cloning site, respectively). pRATDuet derived plasmids carry tetracycline-resistance gene, either single gene or both genes encoding TsuCsx1 and CliCsx15. *E. coli C43 (DE3)* carrying plasmids pCsm1-5 and pCRISPR-TetR or pCRISPR-pUC was transformed with pRATDuet derived plasmids. The cells were recovered for 2 h at 37 °C and serially diluted, before plating onto LB agar supplemented with 100 µ g/ml ampicillin and 50 μg/ml spectinomycin for determination of the recipient cell count. Transformation efficiency was assessed using plates additionally containing 12.5 μg/ml tetracycline. The plasmid immunity was evaluated on plates further supplemented with 0.2 % (w/v) D-lactose and 0.2 % (w/v) L-arabinose, in addition to all three antibiotics. Plates were incubated overnight at 37 °C, then imaged. Technical duplicates of two biological replicates were conducted.

## Data Availability

All experimental materials described in this paper are available from the corresponding author on request. The protein structure co-ordinates and data have been deposited in the Protein Data Bank with deposition codes 9TEU and 9TET.

## Acknowledgements

The authors thank Dr Tracey Gloster for helpful discussion and the University of St Andrews Mass Spectrometry service for technical assistance.

