## Supplementary Material for "The CRISPR ring nuclease Csx15 oligomerises on cyclic nucleotide binding to regulate antiviral defence"

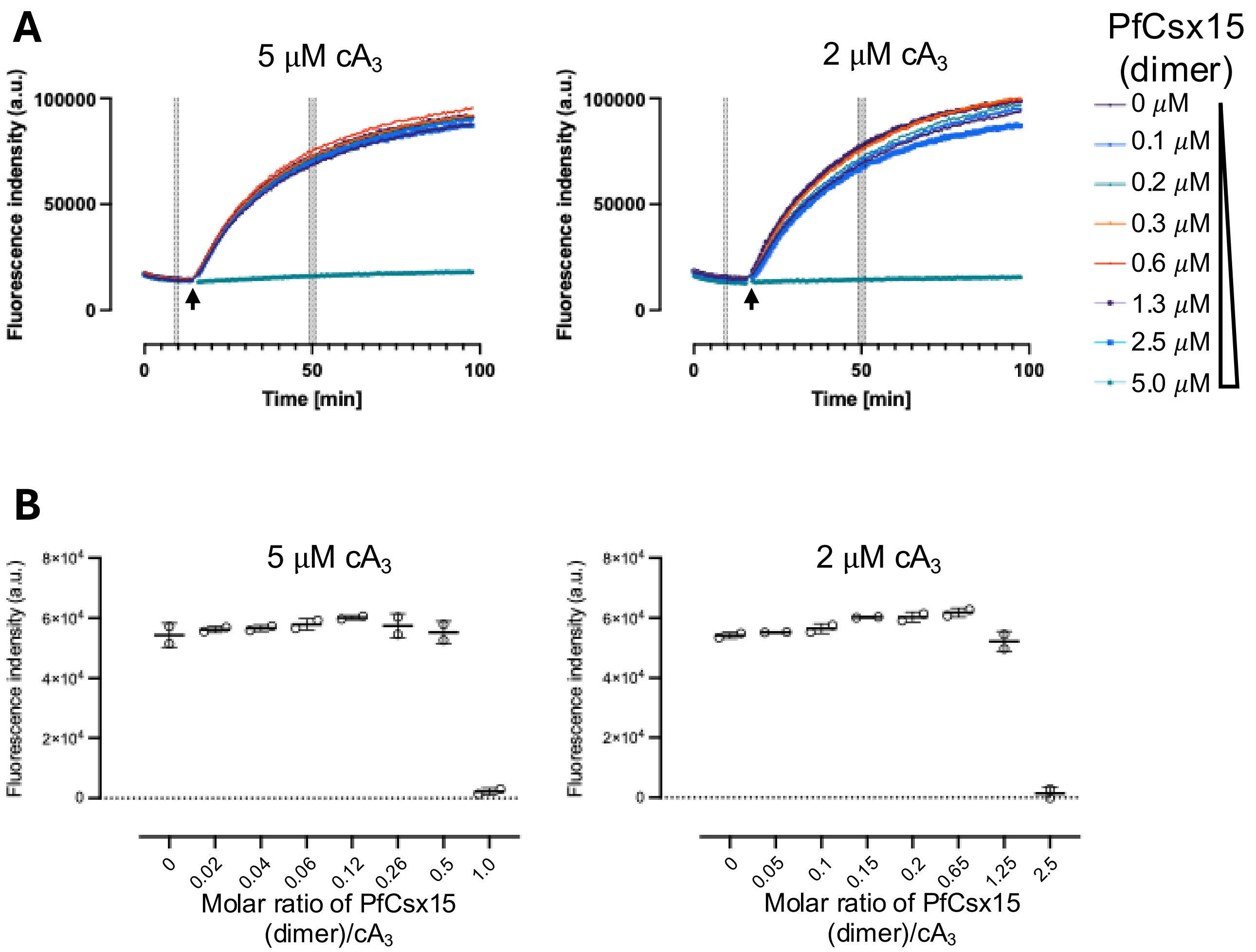

**Supplementary Figure 1: PfCsx15 does not strong inhibit the activated effector VmeNucC**
**A.** Fluorescence signal emitted by dsDNA cleavage when DNases were presented. dsDNA substrates (100 nM) were incubated with the PfCsx15 (0-5 μM dimer) and the $cA_3$ (2 and 5 μM, respectively) at 35°C for 15 min, before adding DNase VmeNucC (250 nM trimer). The fluorescence signal was plotted against time. **B.** Plot of baseline-corrected fluorescence signal in **A** against the ratio of PfCsx15 to $cA_3$. The average fluorescence signal in **A** from 49-51 min was corrected using the average baseline signal from 9-10 min. Data of replicates are presented as mean ± s.d. The DNase activity of NucC was inhibited only when PfCsx15 was present at the highest concentration.

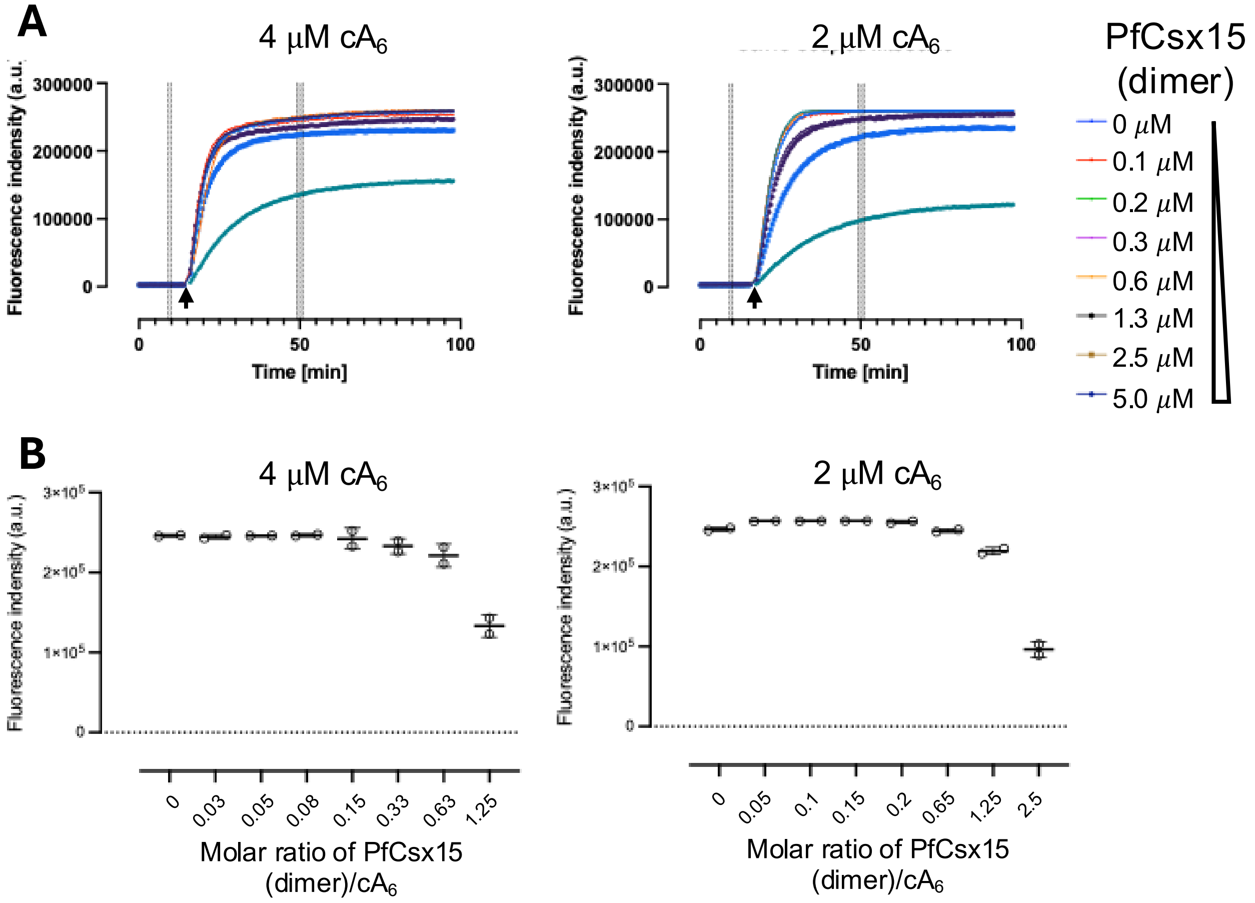

**Supplementary Figure 2. PfCsx15 does not strongly inhibit the $cA_6$-induced effector MtbCsm6**

**A.** Fluorescence signal emitted by RNA cleavage when RNases were presented. RNaseAlert substrates (100 nM) were incubated with the Csx15 (0-5 μM dimer) and the $cA_6$ (2 and 4 μM, respectively) at 35 °C for 15 min, before adding RNase MtbCsm6 (160 nM dimer). The fluorescence signal was plotted against time. **B.** Plot of baseline-corrected fluorescence signal in **A** against the ratio of PfCsx15 to $cA_6$. The average fluorescence signal in **A** from 49-51 min was corrected using the average baseline signal from 9-10 min. Data of replicates are presented as mean ± s.d. The RNase activity of Csm6 was partially inhibited only when PfCsx15 was present at the highest concentration.

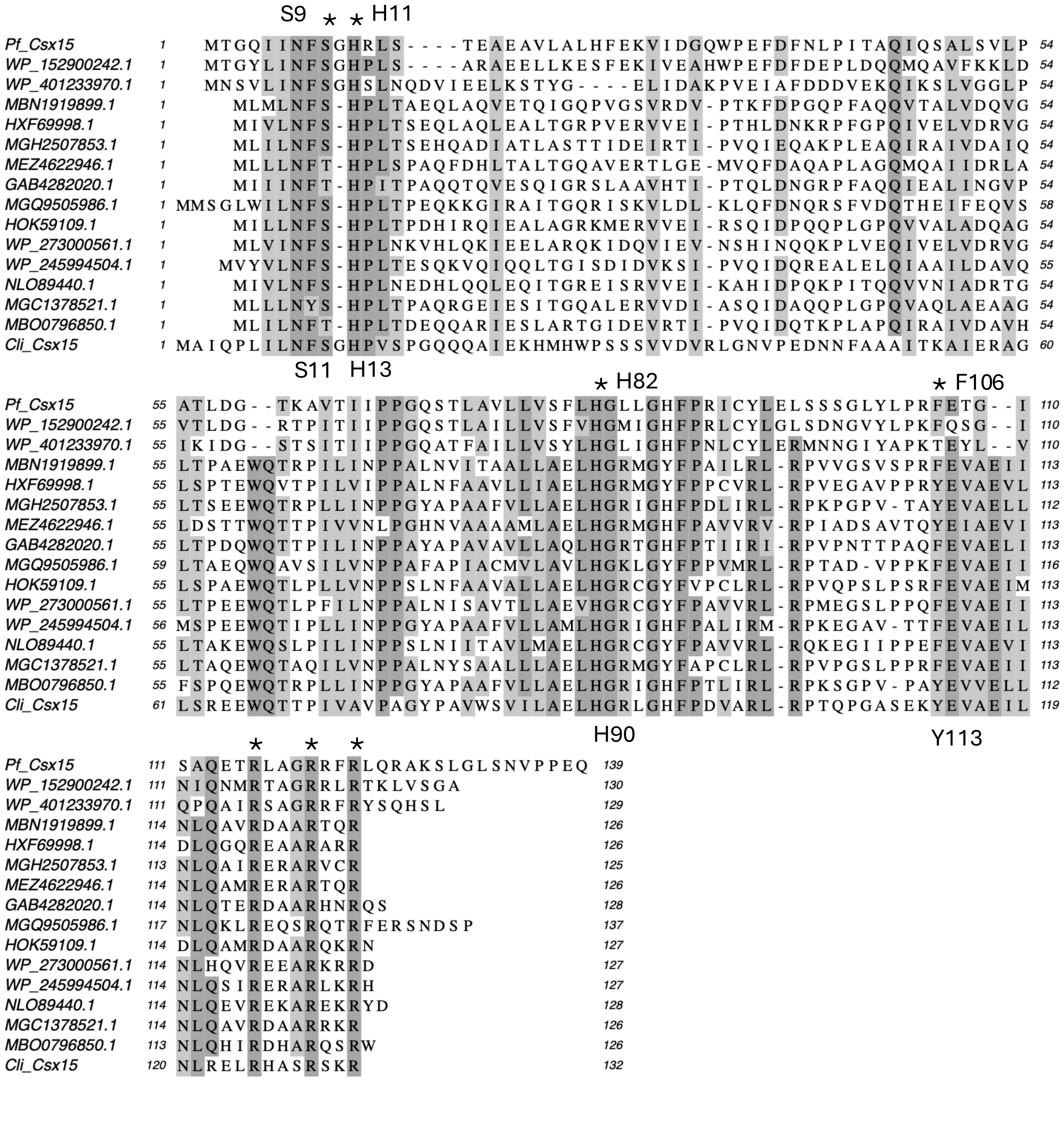

**Supplementary Figure 3. Multiple sequence alignment of a range of Csx15 orthologues.** Conservation is shown by shading and conserved residues mentioned in the text are indicated.

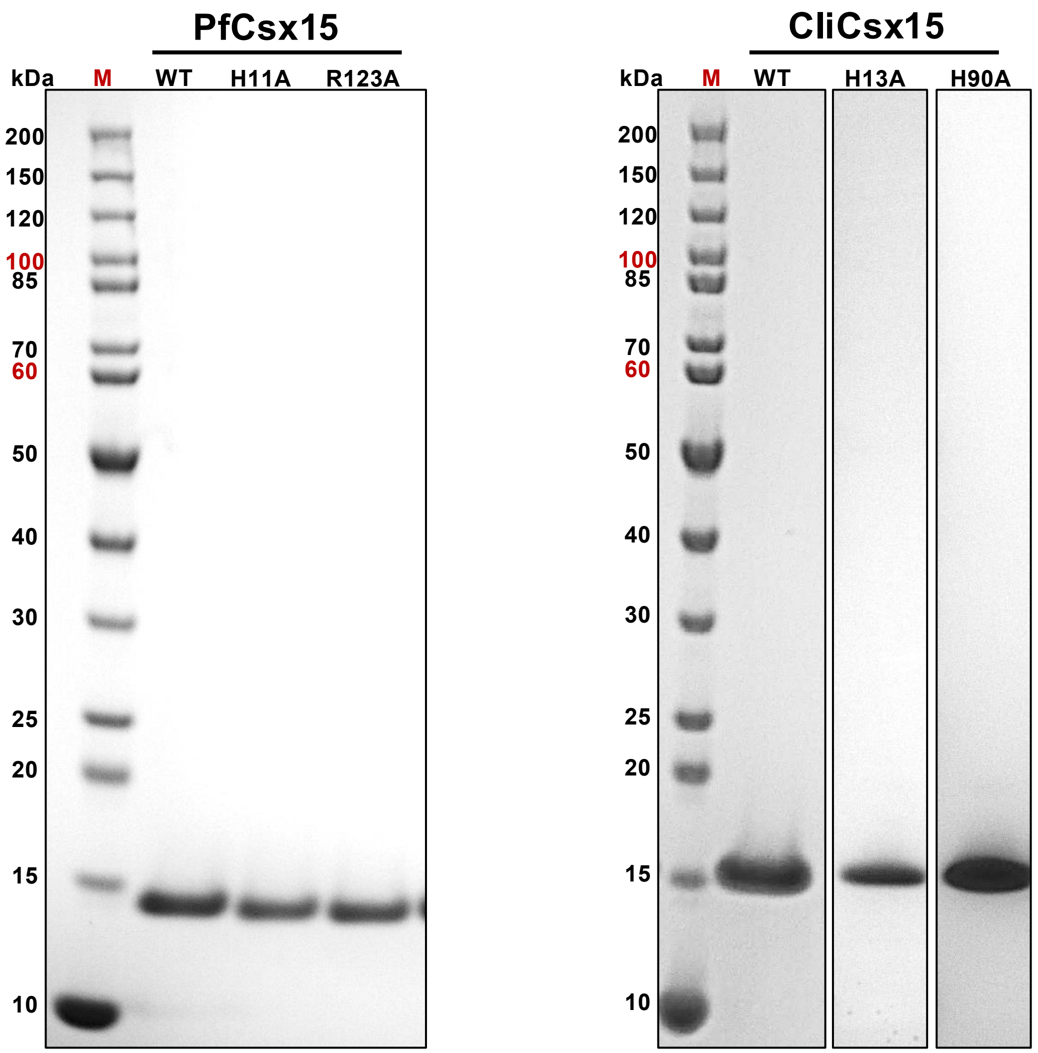

**Supplementary Figure 4. SDS-PAGE analysis of Csx15 wild type and variants.**

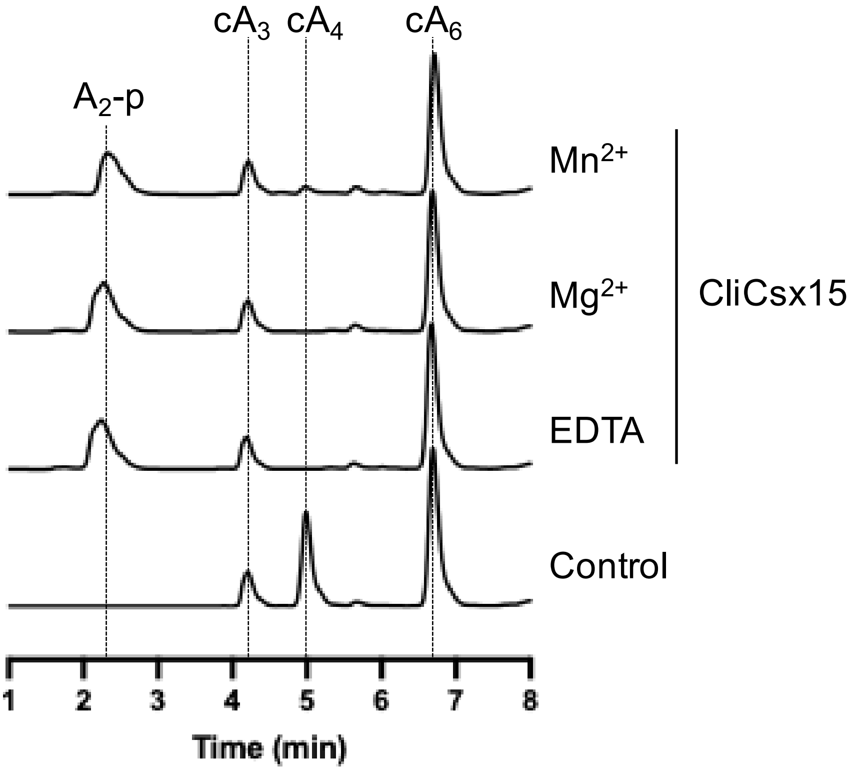

**Supplementary Figure 5. Metal-independent ring nuclease of CliCsx15**

HPLC analysis of CliCsx15 ring nuclease in the presence or absence of metals. The mixture of $cA_3$, $cA_4$ and $cA_6$ was incubated with equal molar CliCsx15 in the presence of 1 mM $MnCl_2$, $MgCl_2$ or EDTA, respectively, for 15 min at 30°C.

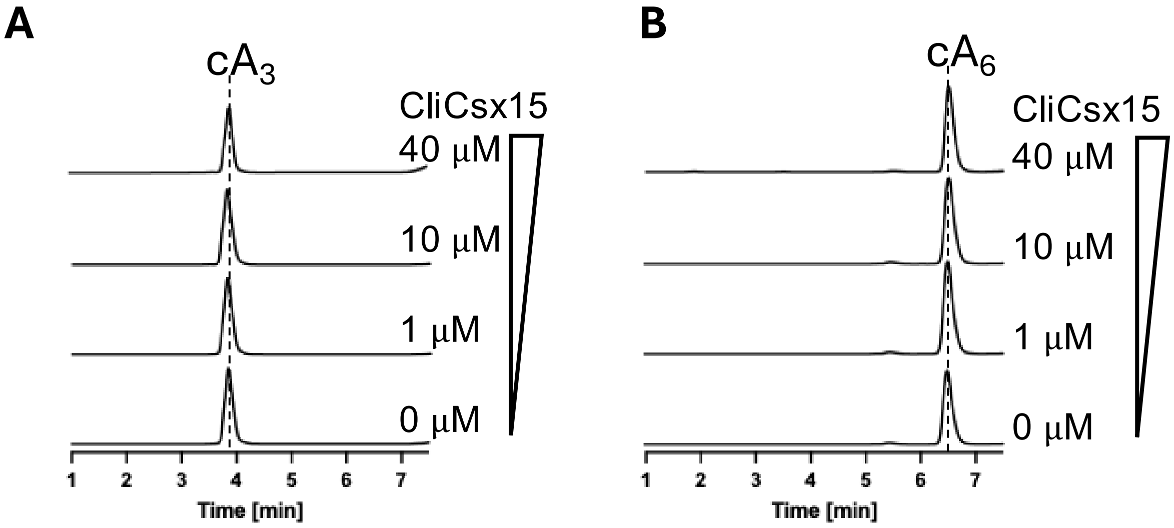

**Supplementary Figure 6. CliCsx15 with $cA_3$ and $cA_6$**

**A.** HPLC analysis of CliCsx15 with $cA_3$ and $cA_6$ in **B**. About 80 µM $cA_3$ and $cA_6$ were incubated with CliCsx15 (0, 1, 10 and 40 ) respectively, for 60 min at 30°C.

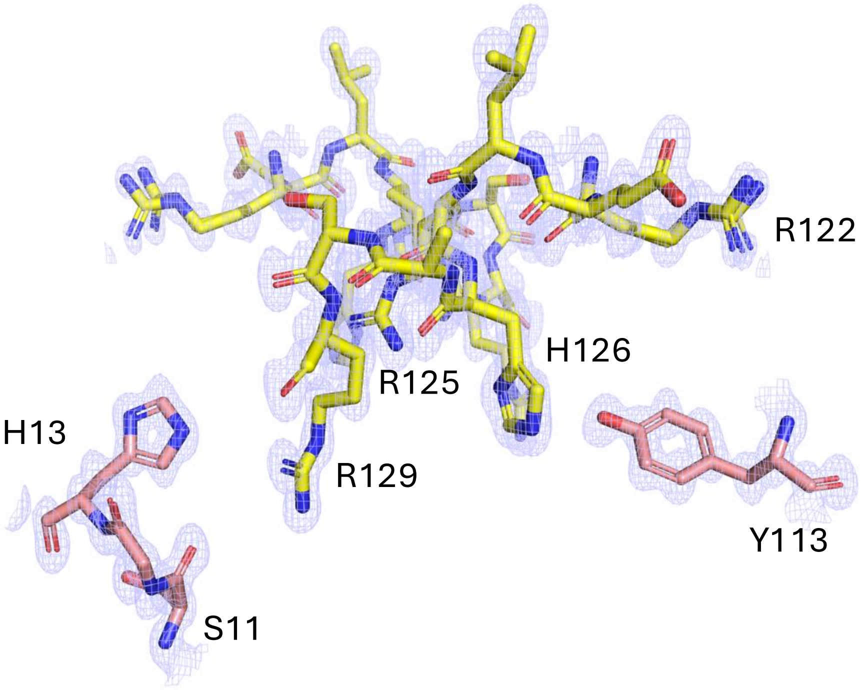

**Supplementary Figure 7. fo-fc electron density map contoured at 1σ across the CliCsx15 dimer interface.** Conserved residues are labelled

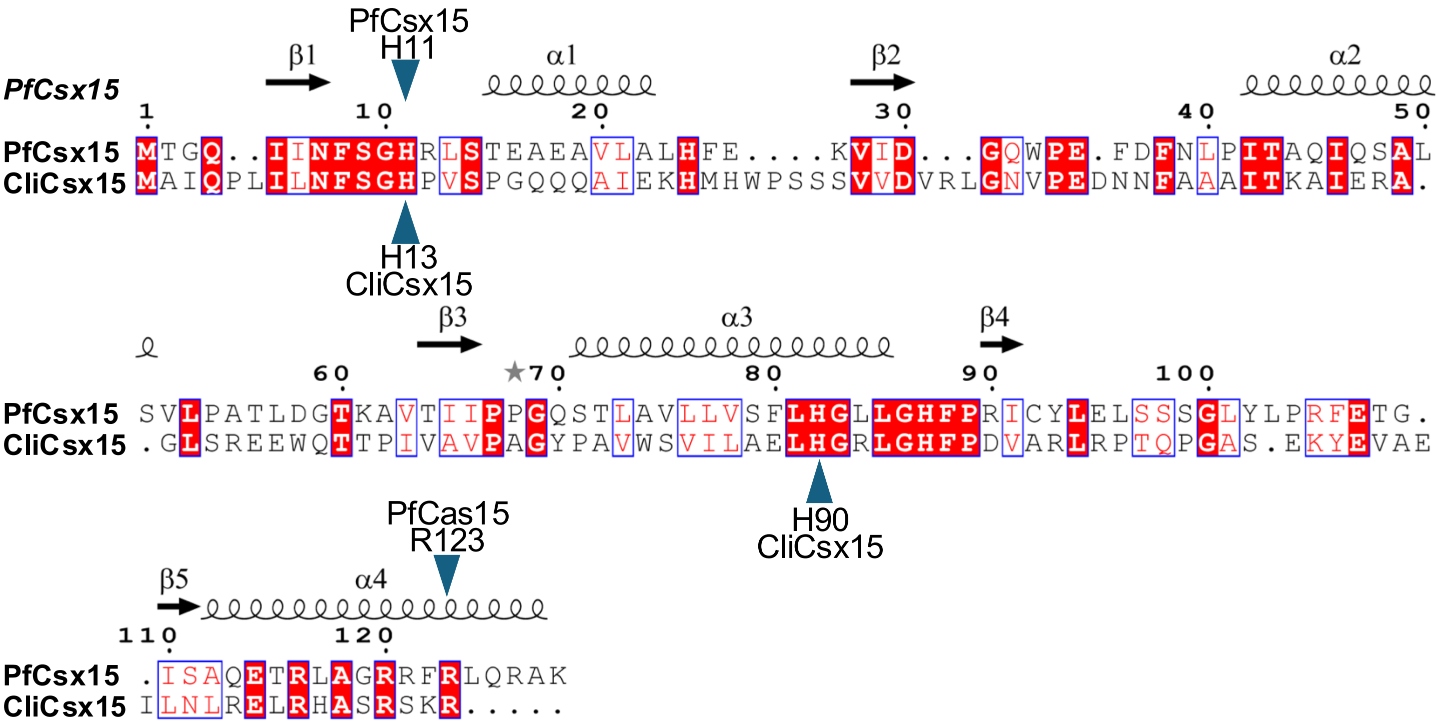

**Supplementary Figure 8. Secondary structure-guided sequence alignment of PfCsx15 and CliCsx15.**

Secondary structure of PfCsx15 is shown above the protein sequence alignment. Residues investigated by site directed mutagenesis are highlighted by blue triangles. Strict conservation is highlighted with the white letter on a red background. Figure was generated using ESPript3 (https://espript.ibcp.fr/ESPript/ESPript/).

### Supplementary Table 1. Data Collection and Refinement Statistics

| | CliCsx15 | PfCsx15 |
|---|---|---|
| **Data processing** | | |
| **Space group** | C 1 2 1 | P62 |
| **Cell dimensions** | | |
| **a, b, c (Å)** | 83.3, 32.5, 57.8 | 89.3, 89.3, 31.4 |
| **α, β, γ (°)** | 90, 132.8, 90 | 90, 90, 120 |
| **Resolution (Å)** | 41.6 – 0.90<br>(0.92 – 0.9) | 77.3 – 1.88<br>(1.91 – 1.88) |
| **$R_{merge}$** | 0.036 (1.320) | 0.135 (3.920) |
| **I/σ(I)** | 16.8 (0.7) | 18.2 (0.5) |
| **Completeness (%)** | 94.6 (55.4) | 99.8 (99.1) |
| **Average redundancy**<br>**$CC_{1/2}$** | 6.2 (3.5)<br>1.000 (0.355) | 40.0 (28.6)<br>1.000 (0.343) |
| **$V_m$ (Å$^3$/Da)** | 1.88 | 2.29 |
| **Solvent (%)** | 34.7 | 46.3 |
| **Refinement** | | |
| **Unique reflections *** | 79628 (2317) | 11875 (582) |
| **$R_{work}$ / $R_{free}$** | 16.6 / 17.6 | 19.7 / 24.7 |
| **Geometric deviations** | | |
| **Bonds (Å) / Angles (°)** | 0.008 / 1.574 | 0.007 / 1.564 |
| **No. atoms (non H)** | | |
| **Protein** | 1100 | 976 |
| **Water**<br>**$PO_4$**<br>**Glycerol** | 202 | 34<br>5<br>6 |
| **B factors (Å$^2$)** | | |
| **Protein** | 13.6 | 60.9 |
| **Water**<br>**$PO_4$**<br>**Glycerol** | 25.0 | 60.1<br>54.2<br>67.0 |
| **Ramachandran**<br>**Favoured / outlier (%)** | 99.2 / 0 | 99.2 / 0 |
| **Molprobity score / centile (%)** | 0.77 / 99 | 1.04 / 100 |
| **Rotamers** | | |
| **Favoured / outlier (%)** | 97.5 / 0 | 96.1 / 0 |
| **PDB Code** | 9TEU | 9TET |

* Values in parentheses are for the highest-resolution shell.

| Name | sequence (5'-3') | Notes |
|---|---|---|
| **Csx15** *Pseudomonas fluorescens* WP_191946534.1 | GCGCCCATGGCACATATGACGGGTCAAATCATCAATTTTTCT<br>GGCCACCGTCTGTCCACTGAAGCGGAGGCCGTACTTGCGCT<br>TCATTTTGAGAAAGTCATTGACGGTCAGTGGCCCGAGTTTGA<br>CTTTAATCTGCCGATCACCGCTCAAATCCAATCAGCGTTGTC<br>AGTTTTGCCCGCAACATTAGACGGCACAAAGGCGGTTACCAT<br>CATTCCCCCTGGCCAGTCGACCCTTGCGGTCTTACTGGTGTC<br>CTTCTTACACGGTTTACTGGGGCATTTCCCGCGTATCTGCTA<br>CCTGGAGCTGTCGTCCTCCGGCCTTTATTTACCCCGCTTTGA<br>GACTGGTATTTCCGCCCAAGAGACGCGTTTGGCTGGCCGTC<br>GTTTCCGCTTACAACGCGCGAAGTCATTGGGCCTGTCGAATG<br>TACCTCCTGAGCAGTGACTCGAGGGATCCCGCG | g-Block |
| **Csx15** *Chlorobaculum limnaeum* WP_069809204.1 | GCGCCCATGGCACATATGGCTATTCAACCGCTTATTTTAAACT<br>TTTCGGGACACCCAGTGTCACCCGGACAGCAACAAGCTATC<br>GAGAAACACATGCATTGGCCGTCCAGTAGCGTGGTGGACGT<br>TCGTCTGGGGAATGTCCCAGAGGATAATAACTTTGCTGCTGC<br>AATCACCAAAGCAATTGAACGCGCTGGGTTGTCACGCGAAGA<br>ATGGCAGACAACGCCAATCGTTGCCGTTCCGGCTGGCTACC<br>CAGCCGTCTGGTCGGTGATTCTGGCCGAGTTACATGGTCGC<br>CTTGGCCATTTTCCGGACGTAGCCCGCTTGCGCCCTACACA<br>GCCTGGGGCGTCCGAGAAGTACGAAGTTGCTGAAATTTTGAA<br>TCTGCGTGAGCTGCGCCACGCATCGCGCTCTAAGCGTTGAC<br>TCGAGGGATCCCGCG | |
| PfCsx15H11A-Fw | CAATTTTTCTGGCGCCCGTCTGTCCACTG | Mutagenesis Primers |
| PfCsx15H111A-Rv | CAGTGGACAGACGGGCGCCAGAAAAATTG | |
| PfCsx15R123A-Fw | GCCGTCGTTTCGCCTTACAACGCGCGAAG | |
| PfCsx15R123A-Rv | CTTCGCGCGTTGTAAGGCGAAACGACGGC | |
| CliCsx15H13A-Fw | CTTATTTTAAACTTTTCGGGAGCCCCAGTGTCAC | |
| CliCsx15H13A-Rv | GCTGTCCGGGTGACACTGGGGCTCCCGAAAAG | |
| CliCsx15H90A-Fw | GATTCTGGCCGAGTTAGCTGGTCGCCTTGGCC | |
| CliCsx15H90A-Rv | GGCCAAGGCGACCAGCTAACTCGGCCAGAATC | |

**Supplementary Table 2. Synthetic genes and mutagenesis primers for Csx15**
